# CwlQ is required for swarming motility but not flagellar assembly in *Bacillus subtilis*

**DOI:** 10.1101/2021.01.13.426625

**Authors:** Sandra Sanchez, Caroline M Dunn, Daniel B. Kearns

## Abstract

Hydrolytic enzymes play an essential role in the remodeling of bacterial peptidoglycan (PG), an extracellular mesh-like structure that retains the membrane in the context of high internal osmotic pressure. Peptidoglycan (PG) integrity must be unfailingly stable to preserve cell integrity but must also be dynamically remodeled for the cell grow, divide and insert macromolecular machines. The flagellum is one such macromolecular machine that transits the PG and the insertion of which is aided by localized activity of a dedicated PG hydrolase in Gram-negative bacteria. To date, there is no known dedicated hydrolase in Gram-positive bacteria for insertion of flagella and here we take a reverse-genetic candidate-gene approach to find that cells mutated for the lytic transglycosylase CwlQ exhibited a severe defect in flagellar dependent swarming motility. We show that CwlQ required its active site to promote swarming, was expressed by the motility sigma factor SigD, and was secreted by the type III secretion system housed inside the flagellum. Nonetheless, cells mutated for CwlQ remained proficient for flagellar biosynthesis even when mutated in combination with four other hydrolases related to motility (LytC, LytD, LytF, and CwlO). The PG hydrolase essential for flagellar synthesis in *B. subtilis*, if any, remains unknown.

## INTRODUCTION

Most bacteria are surrounded by an extracellular cell wall that prevents catastrophic hyper-expansion of the membrane by the high internal osmotic pressure of the cytoplasm. The wall is a semi-elastic macromolecular mesh of peptidoglycan (PG) comprised of long polymers of an *N*-acetyl-glucosamine-*N*-acetyl-muramic acid disaccharide that are cross-linked by amino acid side chains (1,2). While the chemistry of peptidoglycan is relatively invariant, bacteria differ in the organization of PG with respect to overall envelope architecture. In cells with a Gram-negative envelope, the PG is only 1 to 3 layers thick and lays between the plasma and outer cell membrane, whereas cells with a Gram-positive envelope have a much thicker PG wall and lack an outer membrane (3–6). Regardless of the type of envelope, the semi-elastic PG network must be both stable and continuous to maintain cell integrity but dynamically remodeled to allow for cell growth, cell division and the insertion of transenvelope nanomachines.

One nanomachine that is inserted through the peptidoglycan is the propeller-like flagellum that bacteria rotate to swim in liquid or swarm over solid surfaces. Flagella are constructed from over 30 different proteins that are tightly regulated to ensure stoichiometry and sequential assembly (7,8). The first architectural unit of the flagellum to be assembled is the basal body that is inserted in the plasma membrane and houses a dedicated type III secretion system (9,10). Once activated, the type III secretion system secretes the distal components of the flagellum including the structural units of the axle-like rod that is polymerized until it reaches the outer membrane in Gram-negative bacteria, followed by the flexible universal-joint hook (11-16). Hook synthesis terminates when it reaches a particular length, at which point the secretion system transitions to exporting subunits that form the long helical polymer of the filament (17,18). Thus, flagella are constructed from the inside-out and must not only cross all layers of the envelope but freely rotate within them.

The PG is thought to present a structural barrier to flagellar construction at the level of rod (19–21). The rod is the part of the flagellum that spans the PG, and the rod’s diameter of 8-13 nm (22–26) seems incompatibly wide relative to the estimated PG pore size of 2-7 nm (6, 27,28). The first evidence that PG remodeling was required for flagellar assembly came from the observation that mutants in the Gram positive bacterium *B. subtilis* defective in the expression of multiple autolysins (PG hydrolases) were also defective in motility and flagellar biosynthesis (29,30). Later, the role of PG remodeling was further supported in the Gram negative bacteria *Salmonella enterica*, *Rhodobacter sphaeroides*, and *Caulobacter crescentus* when mutants defective in particular PG hydrolases were defective in motility and flagellar assembly (31–35). Remarkably, despite the foundational report, the specific PG hydrolase required for flagellar assembly is not known in *Bacillus subtilis*, and the rod in this organism must penetrate PG that is approximately 50 nm thicker than that of Gram-negative bacteria. Moreover, how the rod transits the peptidoglycan is not known for any Gram-positive bacterium.

*B. subtilis* encodes over 30 annotated PG hydrolases in its genome and here we take a reverse-genetic approach to screen known and putative hydrolases to find genes required for flagellar insertion (36,37). Of the candidates tested, mutation of two PG hydrolases, the vegetative endopeptidase CwlO (38,39) and the poorly understood lytic-transglycosylase CwlQ (40), exhibited a moderate and severe defect in swarming motility, respectively. Seemingly consistent with being a hydrolase involved in flagellar assembly, CwlQ required its active site residue for swarming motility, was expressed by the motility sigma factor SigD and was secreted by the type III secretion system within the flagellum. Inconsistent with being required for flagellar assembly, motility was restored to the *cwlQ* mutant when motility agar concentration was decreased below that of standard swarming conditions and cells mutated for *cwlQ* could both swim and synthesize flagella. Our work suggests that although CwlQ is not required for insertion of the flagella through the PG, it is conditionally required for motility and may play a role in flagella function specifically on harder surface environments. Finally a quintuple mutant disrupting all known SigD-dependent hydrolases and *cwlO* was proficient for flagellar assembly. The PG hydrolase required for flagellar assembly in *B. subtilis*, if any, remains unknown.

## RESULTS

### CwlQ is conditionally required for swarming motility

*B. subtilis* is predicted to encode many peptidoglycan hydrolases, some of which have been biologically and/or biochemically demonstrated to cleave peptidoglycan, and some of which have a predicted function based on sequence homology (36,37) (**Table 1**). A reverse-genetic approach was taken to determine which, if any, of the peptidoglycan hydrolase candidates were required for flagellar assembly in *B. subtilis*. To narrow the pool of candidates, hydrolases and putative hydrolases were excluded if they had been previously tested for flagellar biogenesis (41), if they were expressed only during sporulation or encoded within horizontally-transferred genetic elements (e.g. prophages). The remaining candidate genes were mutated, and the resulting mutants were tested for the flagellar-dependent swarming motility (**Table 1**). Most of the mutants were wild type for swarming behavior and were discarded from further study (**Fig S1**). Cells mutated for either *cwlQ* (**Fig 1A**) or *cwlO* (**Fig 1B**) however, exhibited more severe swarming defects. Moreover, the phenotypes of neither the *cwlQ* nor *cwlO* mutants were due to polar effects on neighboring genes as swarming motility was complemented to wild type when the gene was cloned with 500 bp of upstream DNA (in the case of *cwlQ*) (**Fig 1A**) or expressed from an IPTG-inducible construct (in the case of *cwlO*) (42) (**Fig 1B**) and inserted at an ectopic locus in the respective mutant. We conclude that CwlO and CwlQ are required for swarming motility under standard conditions. CwlO encodes the vegetative endopeptidase required for cell elongation (38,39,42), and we focused our study on the lytic transglycosylase CwlQ as it conferred a more severe swarming defect and its function was poorly-understood (40).

**Figure 1.**
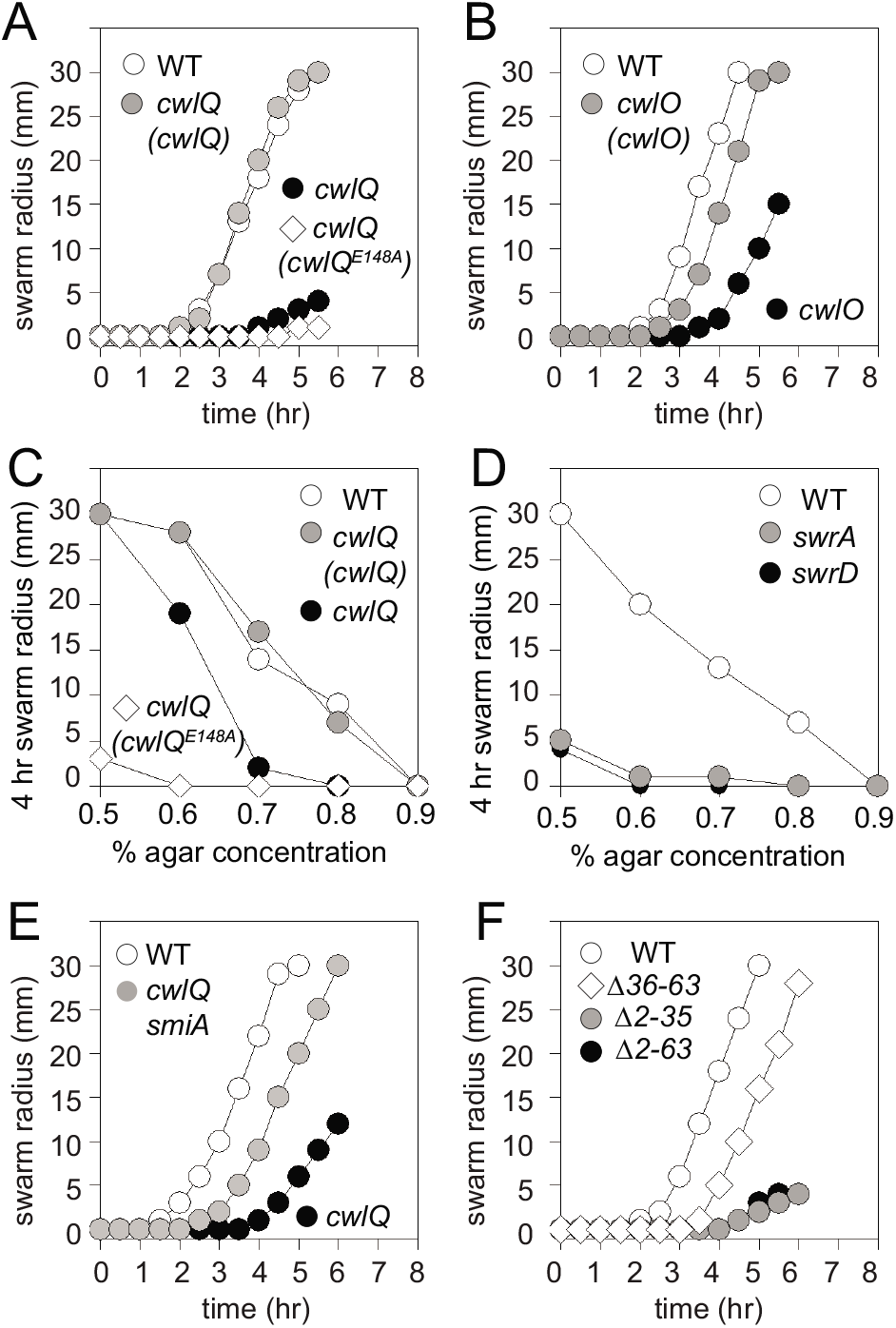
CwlQ is required for swarming motility. A) Quantitative swarm expansion assay of wild type (open circles, DK1042), *cwlQ* (black circles, DK1744), *cwlQ (cwlQ)* (gray circles, DK3586), and *cwlQ (cwlQ^E148A^)* (open diamonds, DK4127). B) Quantitative swarm expansion assay of wild type (open circles, DK1042), cwlO (black circles, DK8462), and cwlO (Pspank-cwlO) in the presence of 1 mM IPTG (gray circles, DK8842). C) Quantitative swarm expansion assay indicating the swarm radius after 4 hours of incubation on media fortified with the indicated amount of agar for the following strains: wild type (open circles, DK1042), *cwlQ* (black circles, DK1744), *cwlQ (cwlQ)* (gray circles, DK3586), and *cwlQ (cwlQ^E148A^)* (open diamonds, DK4127). D) Quantitative swarm expansion assay indicating the swarm radius after 4 hours of incubation on media fortified with the indicated amount of agar for the following strains: wild type (open circles, DK1042), *swrA* (gray circles, DS2415), and *swrD* (black circles, DS6657). E) Quantitative swarm expansion assay of wild type (open circles, DK1042), *cwlQ* (black circles, DK1744), and *cwlQ smiA* (gray circles, DK8018). F) Quantitative swarm expansion assay of cwlQ mutant ectopically complemented with the indicated versions of cwlQ: wild type (open circles, DK3586), D36-63 (open diamonds, DK8472), D2-63 (black circles, DK8470) and D2-35 (gray circles, DK8471). All data points are the average of three replicates.

**Table 1-.** *B. subtilis* candidate PG hydrolase genes

| Gene | Product annotation | Swarming | Excluded |
| --- | --- | --- | --- |
| **Not tested** |  |  |  |
| *blyA* | Muramidase | NT | HTE: SPβ prophage (78) |
| *cwlA* | Muramidase | NT | HTE: Skin element (79) |
| *cwlC* | Muramidase | NT | Sporulation (80,81) |
| *cwlD* | Muramidase | NT | Sporulation ($\sigma^E$, $\sigma^G$) (82) |
| *cwlH* | Muramidase | NT | Sporulation ($\sigma^K$) (83) |
| *cwlP* | Muramidase/endopeptidase | NT | HTE: SPβ prophage (84) |
| *cwlT* | Muramidase/endopeptidase | NT | HTE: ICE element (85,86) |
| *lytH* | Endopeptidase | NT | Sporulation ($\sigma^K$) (87) |
| *spoIID* | Lytic transglycosidase | NT | Sporulation ($\sigma^E$) (88,89) |
| *spoIIP* | Muramidase/endopeptidase | NT | Sporulation ($\sigma^E$) (89,90) |
| *xlyA* | Muramidase | NT | HTE: PBSX prophage (91) |
| *xlyB* | Muramidase (putative) | NT | HTE: PBSX prophage |
| **Wild type swarming** |  |  |  |
| *cwlK* | L,D-endopeptidase | +++ | (92) |
| *lytD* | N-acetylglucosaminidase | +++ | previously tested (41,93,94) |
| *lytE* | Endopeptidase | +++ | (95-97) |
| *lytF* | Endopeptidase | +++ | previously tested (14,98,99) |
| *lytG* | N-acetylglucosaminidase | +++ | (100) |
| *cwlS* | Endopeptidase | +++ | (95,101) |
| *yocH* | Muramidase | +++ | (102) |
| *yqgA* | Wall-associated protein | +++ | (103) |
| *yqgT* | Endopeptidase (putative) | +++ |  |
| *yqil* | Muramidase (putative) | +++ |  |
| *yrvJ* | Muramidase (putative) | +++ |  |
| **Swarming defect** |  |  |  |
| *cwlO* | Endopeptidase | + | (38,39) |
| *cwlQ* | Muramidase/transglycosylase | - | (40) |
| *lytC* | Muramidase | + | previously tested (41,104,105) |

Swarming motility requires flagella and one way in which CwlQ could promote swarming is by remodeling the peptidoglycan to facilitate flagellar assembly (43). To determine whether the *cwlQ* mutant exhibited a flagellar assembly defect, the *cwlQ* gene was mutated in a strain that encoded a variant of the flagellin protein that could be fluorescently labeled with a maleimide dye (Hag^T209C^) (44). After staining, the *cwlQ* mutant was found to be proficient for flagellar filament assembly but appeared to have a qualitative reduction in filament number relative to wild type (**Fig 2A**). Precise counting of filaments in *B. subtilis* is difficult, but filament number can be indirectly assessed by counting the number of flagellar hooks and basal bodies as proxies. Thus, to explore whether the *cwlQ* mutant exhibited a defect in flagellar number, the *cwlQ* gene was mutated in a strain that either encoded a variant of the hook protein that could be fluorescently labeled with a maleimide stain (FlgE^T123C^) or a GFP-fusion to the flagellar basal body protein FliM (12,45). In these backgrounds, the flagellar hooks (**Fig 2B**) and basal bodies (**Fig 2C**) appeared as fluorescent dots, and 3D-structured illumination microscopy was used to count the number of each in both wild type and the *cwlQ* mutant. Quantitative analysis indicated that the there was a subtle but statistically significant reduction (students T-test p value < 0.00003) in the number of flagellar hooks and basal bodies in the *cwlQ* mutant (**Fig 2D, Table S1**). A two-fold increase in flagellar density on surfaces has been shown to be critical for swarming motility in *B. subtilis* (46) and the inability of the *cwlQ* mutant to swarm might be related to the slight reduction in flagellar hook number observed in liquid.

**Figure 2:**
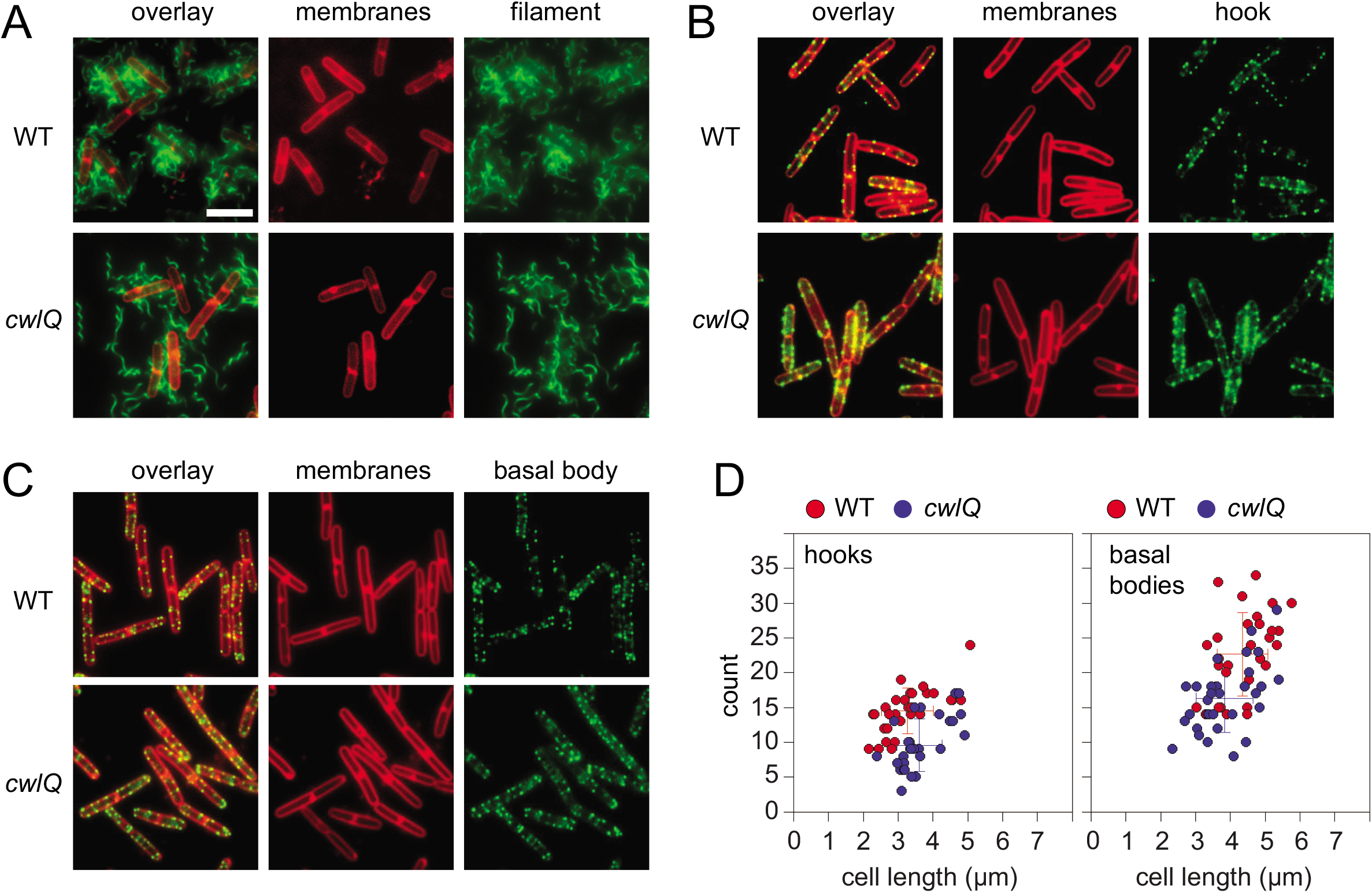
Cells mutated for CwlQ have a slight but statistically significant reduction in the number of flagellar hooks and basal bodies. A) Fluorescence micrographs of cells of the indicated genotype stained for membranes (false colored red) and flagellar filaments (false colored green). The following strains were used to generate this panel: WT (DS1919) and *cwlQ* (DK1770). B) Fluorescence micrographs of cells of the indicated genotype stained for membranes (false colored red) and flagellar hooks (false colored green). The following strains were used to generate this panel: WT (DS7673) and *cwlQ* (DK1771). B) Fluorescence micrographs of cells of the indicated genotype stained for membranes (false colored red) and flagellar basal bodies (FliM-GFP, false colored green). The following strains were used to generate this panel: WT (DS8521) and *cwlQ* (DK1047). D) Scatter plots in which individual wild type (red) and *cwlQ* mutant (blue) cells were measured by OMX 3D-SIM for cell length and the number of flagellar hooks (left) and flagellar basal bodies (right) were counted with Imaris software. Thirty cells were measured per experiment and each cell is represented by a different dot on the graph. Averages and standard deviations are colored according to the data set to which they belong. Raw data is included as supplemental table (**Table S3**).

Another way that the absence of CwlQ might give rise to a swarming defect is if the mutant flagella are defective for rotation. To determine whether the flagella of a *cwlQ* mutant were functional for flagellar rotation, cells were centrally inoculated on LB media fortified with 0.3% agar in which the pores in the agar were sufficiently large to permit swimming motility. As *B. subtilis* preferentially migrates over surfaces, cells were discouraged from surface migration by using a background that was mutated for both surfactant and extracellular polysaccharide biosynthesis (43,47-51). Wild type created a large zone of colonization after 12 hours of incubation whereas a mutant defective in the flagellar filament protein Hag grew as a tight central colony (**Fig 3**). Cells mutated for *cwlQ* produced a zone of colonization similar to that of the wild type (**Fig 3**). Moreover, cells of the *cwlQ* mutant were vigorously motile when grown to exponential phase in liquid media and observed by wet mount microscopy. We conclude that cells mutated for *cwlQ* are not only proficient in flagellar assembly but are also proficient for swimming.

**Figure 3:**
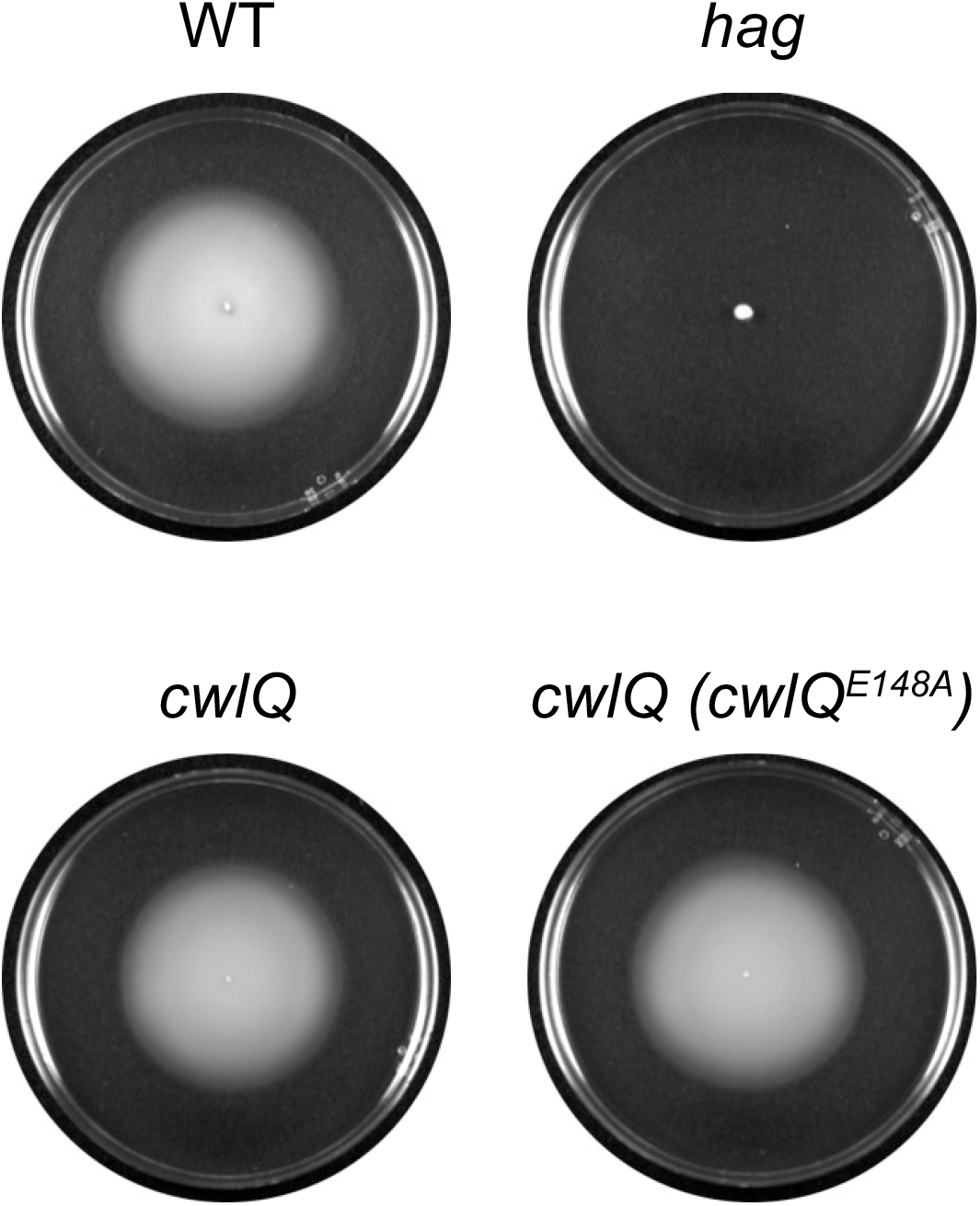
Cells mutated for CwlQ are proficient for swimming motility. LB agar Petri plates fortified with 0.3% agar were centrally inoculated, incubated at 37°C for 12 hours, and filmed against a black background such that zones of colonization appear white and uncololized agar appears black. Each strain contained the indicated alleles plus mutants in *srfAA* and *epsH* to discourage movement across the surface and force cells to swim through the agar. The following strains were used to generate the figure: WT (DK374), *hag* (DK378), *cwlQ* (DK2491), and *cwlQ* (*cwlQ^E148A^*) (DK7051).

As the *cwlQ* mutant exhibited wild type swimming motility in liquid, wild type colonization of 0.3% agar, and only a slight reduction in flagellar hook number, we wondered whether the swarming defect was dependent on the hardness of the agar surface. For the wild type, swarm radius was inversely proportional to agar concentration after 4 hours of incubation and swarming was fully inhibited on media solidified with 0.9% agar (**Fig 1C**). Swarming of the *cwlQ* mutant was fully inhibited after 4 hours at the standard conditions of 0.7% agar, but swarm radius increased with decreasing agar concentration such that the mutant swarmed like the wild type on media solidified with 0.5% agar (**Fig 1C**). Swarming rescue at substandard agar concentrations appeared to be specific to the *cwlQ* mutant as at least two other mutants defective in swarming motility, *swrA* (defective due to reduced flagellar number) (45,52,53) and *swrD* (defective due to reduced flagellar torque) (54), remained non-swarming at all agar concentrations tested (**Fig 1D**). We conclude that the requirement for CwlQ differs from that of other swarming motility mutants as it is conditional and relieved when agar concentrations are reduced below standard conditions.

To determine how CwlQ might promote swarming, we sought to isolate spontaneous suppressors that restored motility to a *cwlQ* mutant upon prolonged incubation on a swarm agar plate. Unlike regulatory mutants defective in swarming (14,41,50,54,55), no spontaneous swarming-proficient suppressor ever emerged as a flare from the non-motile colony of the *cwlQ* mutant, even after 48 hours of incubation. To directly test the hypothesis that flagellar number was limiting in the *cwlQ* mutant, a double mutant was generated that was simultaneously defective in *cwlQ* and *smiA* encoding SmiA, a specific adaptor protein for the regulatory proteolysis of SwrA (46). Flagellar number and swarming increases when *smiA* is mutated (46), and the *cwlQ smiA* double mutant exhibited enhanced swarming motility relative to the *cwlQ* mutant alone (**Fig 1E**). We infer that the swarming defect in the absence of CwlQ is likely structural rather than regulatory, because like mutants defective in flagellar structure, spontaneous suppressor mutants could not be isolated. We further infer that CwlQ is modestly defective in flagellar number, and swarming motility can be improved either by reducing surface hardness or by increasing flagellar number though mutation of SmiA.

### CwlQ is secreted and swarming requires the CwlQ active site

To further explore the mechanism of CwlQ, the CwlQ primary sequence was analyzed. CwlQ is predicted to have two domains: an N-terminal domain of unknown function and a C-terminal lytic transglycosylase domain previously shown to require a conserved glutamate for peptidoglycan hydrolase activity (40,56,57) (**Fig 4A**). To determine whether lytic transglycosylase activity was required for swarming, the conserved glutamate active site residue E148 was mutated to an alanine (*cwlQ^E148A^*) in the complementation construct and inserted at an ectopic locus (*amyE::P_cwlQ_-cwlQ^E148A^*) in a *cwlQ* mutant background. Introduction of the active site mutant allele conferred a defect in swarming motility that was somewhat more severe than the *cwlQ* mutant alone (**Fig 1B**). Consistent with an enhanced defect, the strain that expressed the *cwlQ^E148A^* allele exhibited reduced swarm expansion rate relative to the *cwlQ* null mutant at all agar concentrations tested (**Fig 1C**). Finally, the defect appeared to be specific for swarming as the *cwlQ^E148A^* mutant exhibited swimming motility like the wild type (**Fig 3**). We conclude that CwlQ requires the lytic transglycosylase active site to promote swarming motility and that the presence of CwlQ may become inhibitory when the active site is abrogated.

**Figure 4.**
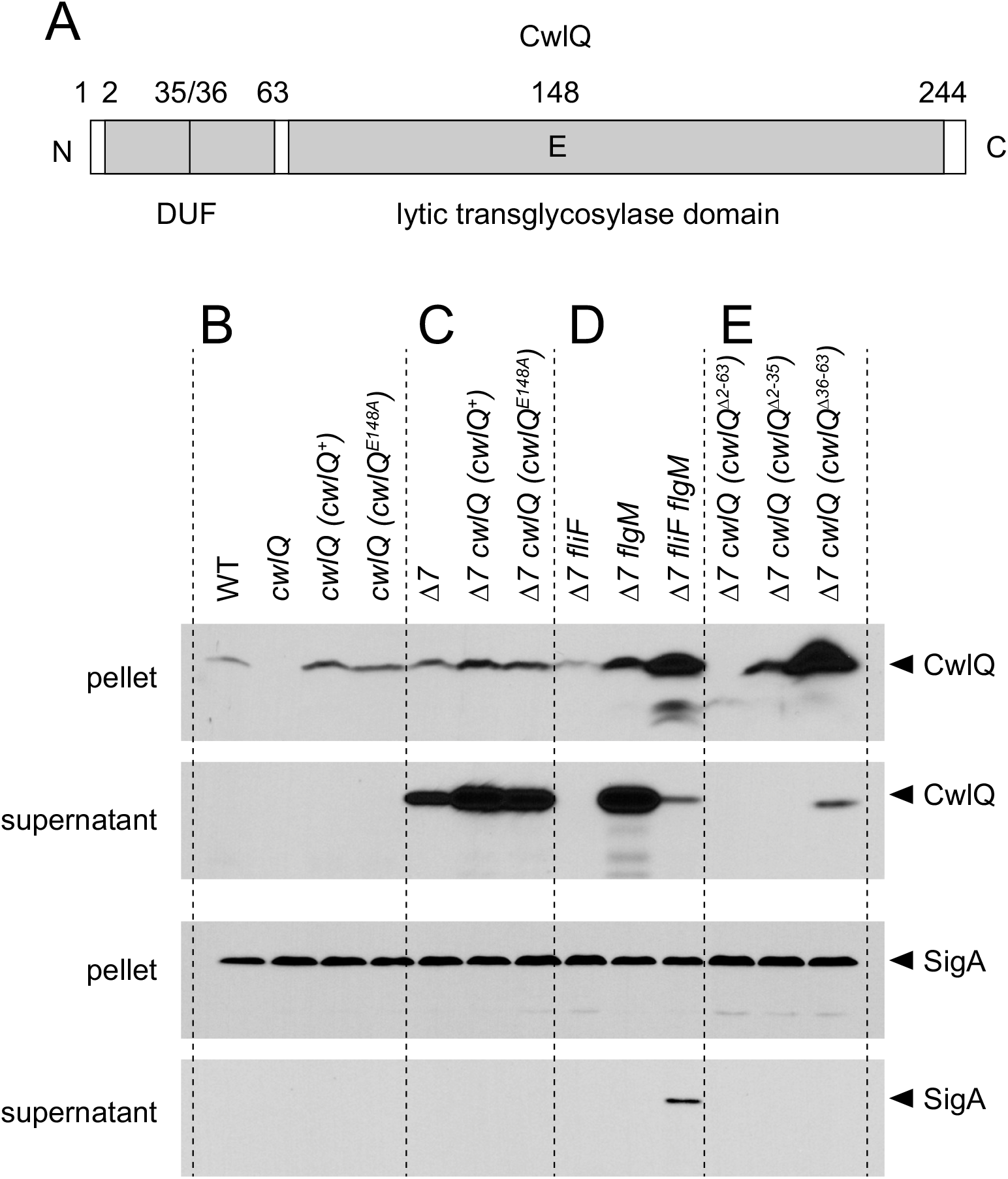
CwlQ is secreted by the flagellar type III secretion system and destroyed by extracellular proteases. A) Cartoon diagram of the 244 amino acid CwlQ protein primary sequence from N-terminus (left) to C-terminus (right). Important amino acid residue numbers are indicated above the diagram and the domains of unknown function (DUF) and lytic transglycosylase domains are indicated below. B-E) Western blot analysis of cell pellets and TCA precipitated supernatants resolved by SDS-PAGE and probed either with anti-CwlQ or anti-SigA antibodies. Panel B) WT (DK1042), *cwlQ* (DK1744), *cwlQ* (*cwlQ*) (DK3586), and *cwlQ* (*cwlQ^E148A^*) (DK4127). Panel C) *Δ7* (DS6329), *Δ7 cwlQ (cwlQ)* (DK8697), and *Δ7 cwlQ* (*cwlQ^E148A^*) (DK8698). Panel D) *Δ7 fliF* (DS6871), *Δ7 flgM* (DS7160), *Δ7 fliF flgM* (DK5150). Panel E) *cwlQ (cwlQ^Δ2-63^)* (DK8699), *cwlQ (cwlQ^Δ2-35^*) (DK8700), and *cwlQ (cwlQ^Δ36-63^)* (DK8701).

As lytic transglycosylases operate on the extracellular substrate peptidoglycan, we hypothesized that CwlQ was secreted from the cytoplasm. In order to determine if CwlQ was secreted, cell lysates and TCA-precipitated supernatants were probed with anti-CwlQ and anti-SigA antibody in Western blot analysis (**Fig 4B**). The cytoplasmic housekeeping sigma factor SigA was used both as a loading control for the cytoplasmic fraction, and its absence in the supernatant indicated that protein release by spontaneous cell lysis was likely minimal (**Fig 4B**). CwlQ was present in cell lysates of wild type, absent in the *cwlQ* mutant, and was restored in the CwlQ complementation strain (**Fig 4B**). Moreover, CwlQ protein was also detected when CwlQ^E148A^ was expressed in an otherwise *cwlQ* mutant background suggesting that the active site mutant was not defective due to inherent protein instability (**Fig 4B**). There was no indication of extracellular CwlQ in the strains tested (**Fig 4B**).

A failure to detect extracellular CwlQ could either indicate that CwlQ was not secreted and functioned in the cytoplasm, or that it was secreted and subsequently degraded by extracellular proteases as has been shown for other flagellar proteins in *B. subtilis* (16,58). To determine whether secreted proteases contributed to extracellular CwlQ degradation, pellets and TCA-precipitated supernatants were harvested, resolved, and subjected to Western blot analysis in a variety of strains deleted for seven extracellular proteases (Δ7) (58). The CwlQ protein was found in the both the pellet and TCA-precipitated supernatant fraction of the otherwise wild type, the *cwlQ* mutant complemented with wild type allele, and the *cwlQ* mutated complemented with the CwlQ^E148A^ allele when the seven extracellular proteases were absent (**Fig 4C**). We conclude that CwlQ is a secreted protein that cannot normally be detected in the supernatant due to extracellular degradation by one or more of the proteases secreted by *B. subtilis*.

The primary sequence of CwlQ does not encode signal sequences consistent with either SEC-dependent or TAT-dependent secretion (59–62). One way in which CwlQ could be secreted in manner independent of a known signal sequence is if CwlQ was secreted by the type III secretion system that resides at the core of the flagellum (63–65). To determine whether CwlQ was secreted by the flagellar type III system, cells were mutated for the basal body protein FliF, a protein previously shown to be essential for flagellar-mediated secretion (58). CwlQ protein was detected in the cytoplasm but not the supernatant of the *fliF* Δ7 mutant (**Fig 4D**). The absence of CwlQ from the supernatant could be consistent with a failure of secretion, but many genes that are required for flagellar motility are under the regulation of the alternative sigma factor SigD, and SigD activity is inhibited by the anti-sigma factor FlgM when *fliF* is mutated (58,66-69). Moreover, the total amount of CwlQ protein appeared to be reduced in the *fliF* Δ7 mutant perhaps consistent with an expression defect (**Fig 4D**). Thus, if *cwlQ* was expressed as part of the flagellar regulon, it would be difficult to distinguish whether the absence of CwlQ protein in the supernatant in a *fliF* mutant was either due to a failure of secretion or a failure of expression, or both.

To determine whether the expression of *cwlQ* was impaired in a *fliF* mutant, a transcriptional reporter construct was generated in which the *cwlQ* promoter region (*P_cwlQ_*) was fused to the *lacZ* gene encoding β-galactosidase and inserted at an ectopic site (*amyE::P_cwlQ_-lacZ*). Mutation of *fliF* reduced the expression of the *P_cwlQ_-lacZ* reporter 10-fold (**Fig 5**). Consistent with the *fliF* defect, expression of *cwlQ* was found to be SigD-dependent and *cwlQ* expression was abolished when SigD was mutated (**Fig 5**). Moreover, mutation of *flgM* increased *P_cwlQ_* expression above that of the wild type, and *P_cwlQ_* expression was restored to the *fliF* mutant when *flgM* was also disrupted (**Fig 5**). We conclude that *cwlQ* is a SigD-dependent gene and the lack of extracellular CwlQ in the absence of FliF may have been due, at least in part, to a protein expression failure. To determine whether CwlQ was secreted in the *fliF* mutant without the confounding expression defect, Western blot analyses were conducted on cells mutated for *flgM* and a *fliF flgM* double mutant in the Δ7 background. CwlQ secretion was dramatically reduced in the *fliF flgM* double mutant background relative to the *flgM* single mutant alone (**Fig 4D**). We note that the cytoplasmic control SigA protein also appeared in the supernatant of the *fliF flgM* double mutant and thus the small amount of CwlQ that appeared to be secreted may have been spuriously released by cellular lysis. We conclude that CwlQ is primarily, and likely exclusively, secreted in a manner that depends on FliF and the flagellar type III secretion system.

**Figure 5.**
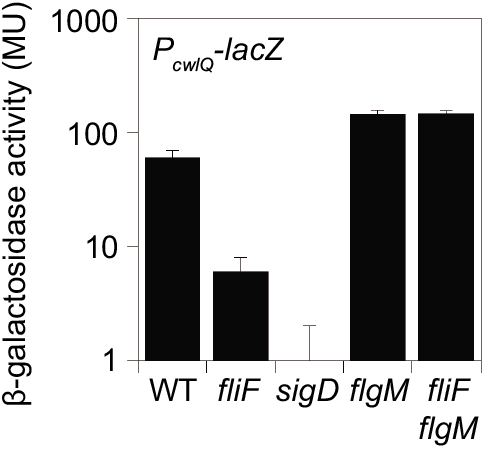
The cwlQ gene is expressed by RNA polymerase and the alternative sigma factor, SigD. β-galactosidase activities of a transcriptional fusion of the promoter of *cwlQ* to the *lacZ* gene encoding β-galactosidase (*P_cwlQ_-lacZ*) in the indicated genetic backgrounds and expressed in Miller units (MU). Each bar is the average of three replicates and standard deviations are provided. The following strains were used to generate the figure: WT (DK2185), *fliF* (DK8864), *sigD* (DK2207), *flgM* (DK2347), and *fliF flgM* (DK8665).

The signal sequence that directs proteins to be secreted by type III secretion system is poorly-understood but appears to be contained within the N-terminus of a secreted protein (63-65). To determine whether the N-terminal domain of CwlQ was required for secretion, three separate in-frame markerless deletions were generated that separately deleted amino acids 2-63 (deleting the entire N-terminus), and 2-35 and 36-63 (deleting the first and second halves of the N-terminus respectively) in the *cwlQ* complementation construct (**Fig 4A**). CwlQ^Δ2-63^ and CwlQ^Δ2-35^ both displayed a defect in swarming motility comparable to that of the *cwlQ* null mutation when introduced to a strain deleted for the native copy of *cwlQ*, but CwlQ^Δ36-63^ swarmed like wild type albeit with an extended lag period (**Fig 1F**). When CwlQ^Δ2-63^ was expressed in a Δ7 strain deleted for extracellular proteases, no protein was detected suggesting that the deletion of the entire N-terminal domain caused severe defects in protein stability (**Fig 4E**). CwlQ^Δ2-35^ was detected in the cell pellet but not the supernatant suggesting that the N-terminus of CwlQ was required for secretion (**Fig 4E**). Finally CwlQ^Δ36-63^ hyper-accumulated in the cytoplasm with a reduced level of secretion that might account for the prolonged rescued of swarming to the *cwlQ* mutant (**Fig 4E**). We conclude that the N-terminus of CwlQ is important for its secretion and protein stability.

### CwlQ and CwlO are not synergistically required for flagellar assembly

Cells mutated for *cwlQ* alone exhibited a conditional defect in swarming motility and were proficient in flagellar assembly. One reason that a peptidoglycan hydrolase mutant might fail to have flagellar assembly defect is if other peptidoglycan hydrolases with which it is co-expressed were redundant in supporting the activity. Since *cwlQ* is a fourth secreted peptidoglycan hydrolase expressed as part of the SigD-regulon (41), a quadruple mutant was generated defective in *cwlQ lytC lytD lytF* in a background that expressed a version of the flagellar filament that could be fluorescently labeled. The quadruple mutant was still proficient for flagellar synthesis (Fig 6A). Next, a *cwlO* mutation was introduced to the *cwlQ lytC lytD lytF* quadruple mutant and the resulting quintuple mutant showed increased loss of cell integrity (by an increase in frequency of cytoplasmic staining with the maleimide dye), defects in cell shape, and reduced flagellar filament number (Fig 6B). The *cwlO* mutant alone did not exhibit a severe reduction in flagellar filament number (Fig 6C), and neither did a *cwlO cwlQ* (Fig 6D) nor a *cwlO lytC* double mutant (Fig 6E). We conclude that neither CwlQ nor CwlO are essential for flagellar assembly and that mutation of as many as five peptidoglycan hydrolases is required to reduce, but not abolish, flagellar synthesis and/or retention.

**Figure 6:**
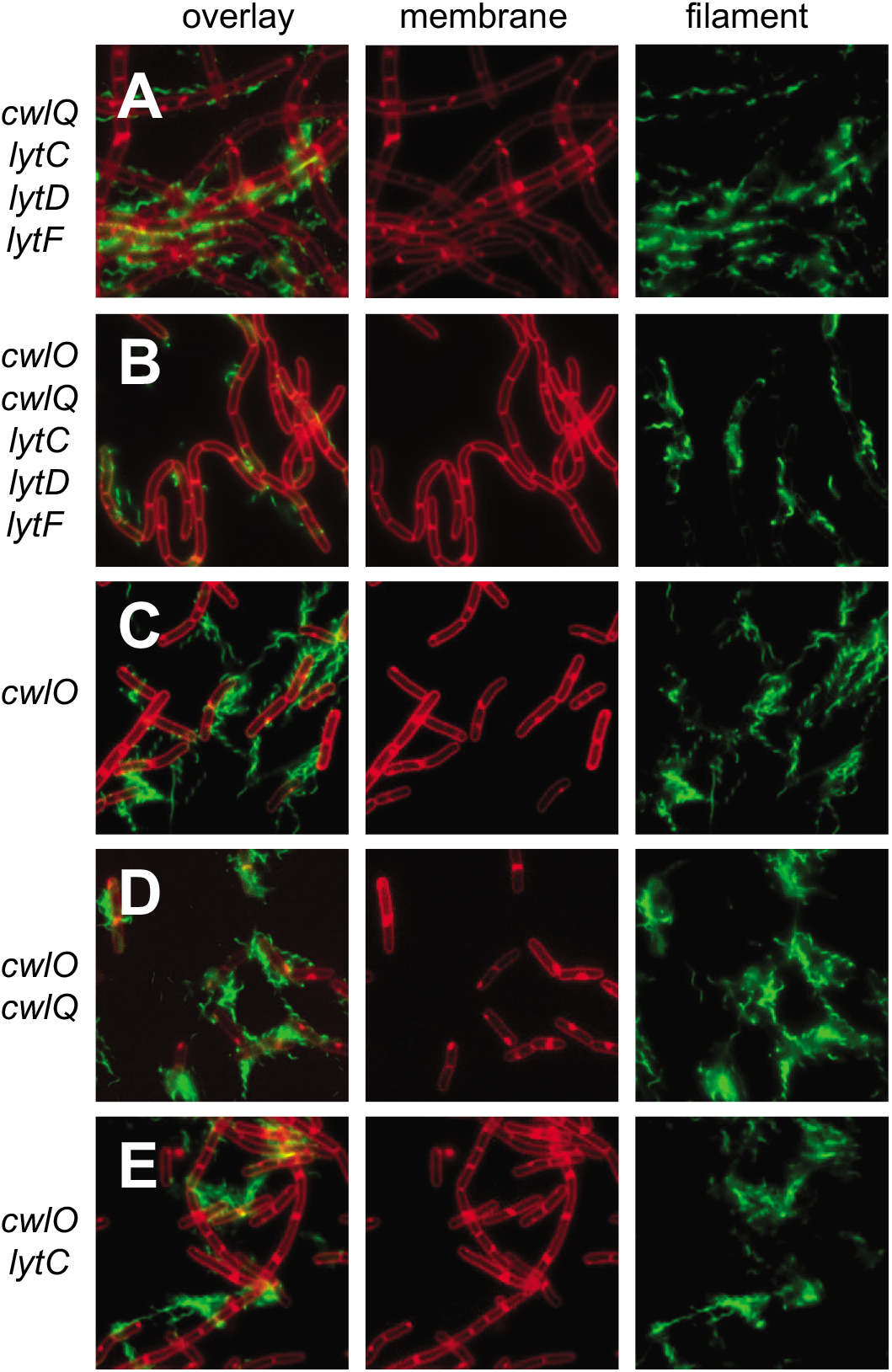
A strain simultaneously mutated for CwlQ, CwlO, and three other PG hydrolases is proficient for flagellar biosynthesis. Fluorescence micrographs of cells of the indicated genotype stained for membranes (false colored red) and flagellar filaments (false colored green). The following strains were used to generate this panel: *cwlQ lytC lytD lytF* (DK7570), *cwlQ cwlO lytC lytD lytF* (DK8815) *cwlO* (DK8816), *cwlQ cwlO* (DK8783), and *cwlO lytC* (DK8787).

## Discussion

Flagella are elaborate molecular nanomachines, which when assembled span the plasma membrane and all layers of the cell envelope including the peptidoglycan (PG) wall. The macromolecular structure of PG is porous with pores approximately 2-7 nm in diameter (22-26), and peptidoglycan remodeling by dedicated hydrolases is thought necessary to allow insertion of the 8-13 nm flagellar rod (6,27,28). Gram negative bacteria often encode a PG hydrolase that has a second function in flagellar rod polymerization such that as the rod extends towards the PG, the hydrolase degrades the wall and permits rod passage (31,33,34). Gram positive bacteria are surrounded by PG many layers thick but to date, no PG hydrolase has been shown to be required for flagellar assembly. Here we took a reverse-genetic, candidate-gene approach to identify PG hydrolases required for flagellar assembly in *B. subtilis* and found that mutation of two hydrolases CwlO and CwlQ resulted in defects in swarming motility. Neither mutant was defective in flagellar assembly however, either when mutated singly or when mutated in combination with four other PG hydrolases. To better understand the role of PG hydrolases in promoting motility, we focused on the less well-understood CwlQ.

Cells mutated for CwlQ were defective for swarming motility under standard conditions but the reason for the defect is unclear. We suspect that CwlQ plays a structural role in swarming as, unlike the case of mutants defective in swarming for regulatory reasons, no spontaneous suppressor mutants were isolated that restored swarming to the cwlQ mutant. Perhaps consistent with a structural role, there appeared to be a qualitative reduction in flagellar filaments and we often saw evidence of filaments dissociated from the cell body in fluorescent micrographs, even in liquid-grown culture, of the *cwlQ* mutant. Perhaps the absence of CwlQ creates a local environment in the peptidoglycan that promotes fracture/instability of the rod. Additionally, the *cwlQ* mutant exhibited a statistically significant reduction in both the number of flagellar basal bodies and hooks, and swarming was somewhat improved by additional mutation of SmiA, a protein that restricts flagellar number. How or why basal body number might be reduced in the absence of CwlQ is unclear however, as basal body assembly is thought to precede interaction with the peptidoglycan. Finally, swarming could be restored to the *cwlQ* mutant simply by reducing the agar concentration of the surface. Previous work indicated that the transition to swarming motility in *B. subtilis* requires that cells exceed a threshold flagellar density (46) and cells mutated for CwlQ, when introduced to a surface, may be below that level for 0.7% agar, but above a reduced level needed for softer agars.

While the mechanism by which CwlQ promotes swarming is unknown, here we make a number of observations that connects the function of CwlQ to flagellar structure and/or activity. First, a conserved and biochemically-determined active site residue for CwlQ lytic transglycosylase activity (40) was required to promote swarming and when mutated caused even more severe defect than deletion of entire gene, perhaps suggesting that unproductive binding of CwlQ to PG becomes inhibitory. Second, CwlQ was found to be part of the flagellar regulon under strict control of the flagellar sigma factor SigD, and was thus co-expressed with, among other proteins, the structural subunits of the distal rod and flagellar filament. Third, CwlQ was secreted in a manner dependent on the type III secretion system housed within the flagellum and its export was directed by information encoded within the poorly-conserved N-terminal domain. Thus, CwlQ promoted swarming motility as a lytic transglycosylase, was co-expressed with flagellar structural subunits and was secreted by the flagellum. Every observation above makes CwlQ seem an ideal candidate to be a PG remodeling hydrolase for flagellar assembly and yet, flagella were synthesized in its absence.

One reason, often invoked, for why mutation of a PG hydrolase fails to confer phenotype is the idea of redundancy, that the genome encodes another peptidoglycan hydrolase with redundant activity or that otherwise compensates for the absence. At the most basic level, all PG hydrolases are redundant as they all operate on the same substrate, peptidoglycan, but what distinguishes them is how their activity is restricted in both space and time. Possible candidates for redundant activity with CwlQ are the LytC, LytD, and LytF hydrolases with which it is co-expressed and the CwlO PG hydrolase that promotes cell elongation and is required for full swarming motility. Simultaneous mutation of all five hydrolases reduced, but failed to abolish, flagellar production. The quintuple mutant also showed signs of reduced cellular integrity in the form of frequently misshapen and lysed cells, and thus the effect of the quintuple deletion may be less specific for flagellar synthesis and more an indication of generalized envelope damage. Whatever the case, the identity of the specific PG hydrolase dedicated to flagellar transit of the wall during assembly in *B. subtilis*, remains unknown.

## MATERIALS AND METHODS

### Strains and growth conditions

*B. subtilis* strains were grown in Luria-Bertani (LB) (10 g tryptone, 5 g yeast extract, 5 g NaCl per L) broth or on LB plates fortified with 1.5% Bacto agar at 37°C. When appropriate, antibiotics were included at the following concentrations: 10 μg/ml tetracycline, 100 μg/ml spectinomycin, 5 μg/ml chloramphenicol, 5 μg/ml kanamycin, and 1 μg/ml erythromycin plus 25 μg/ml lincomycin (*mls*). Isopropyl β-D-thiogalactopyranoside (IPTG, Sigma) was added to the medium at the indicated concentration when appropriate.

### Strain construction

All constructs were either first introduced by transformation by natural competence into DK1042 (a competent derivative of strain 3610) (70), or transformed into the domesticated strain PY79 and transduced in 3610 using SPP1-mediated generalized phage transduction (71). All strains used in this study are listed in Table 2. All primers used in this study are listed in Table S2. All plasmids used in this study are listed in Table S3.

**Table 2:**
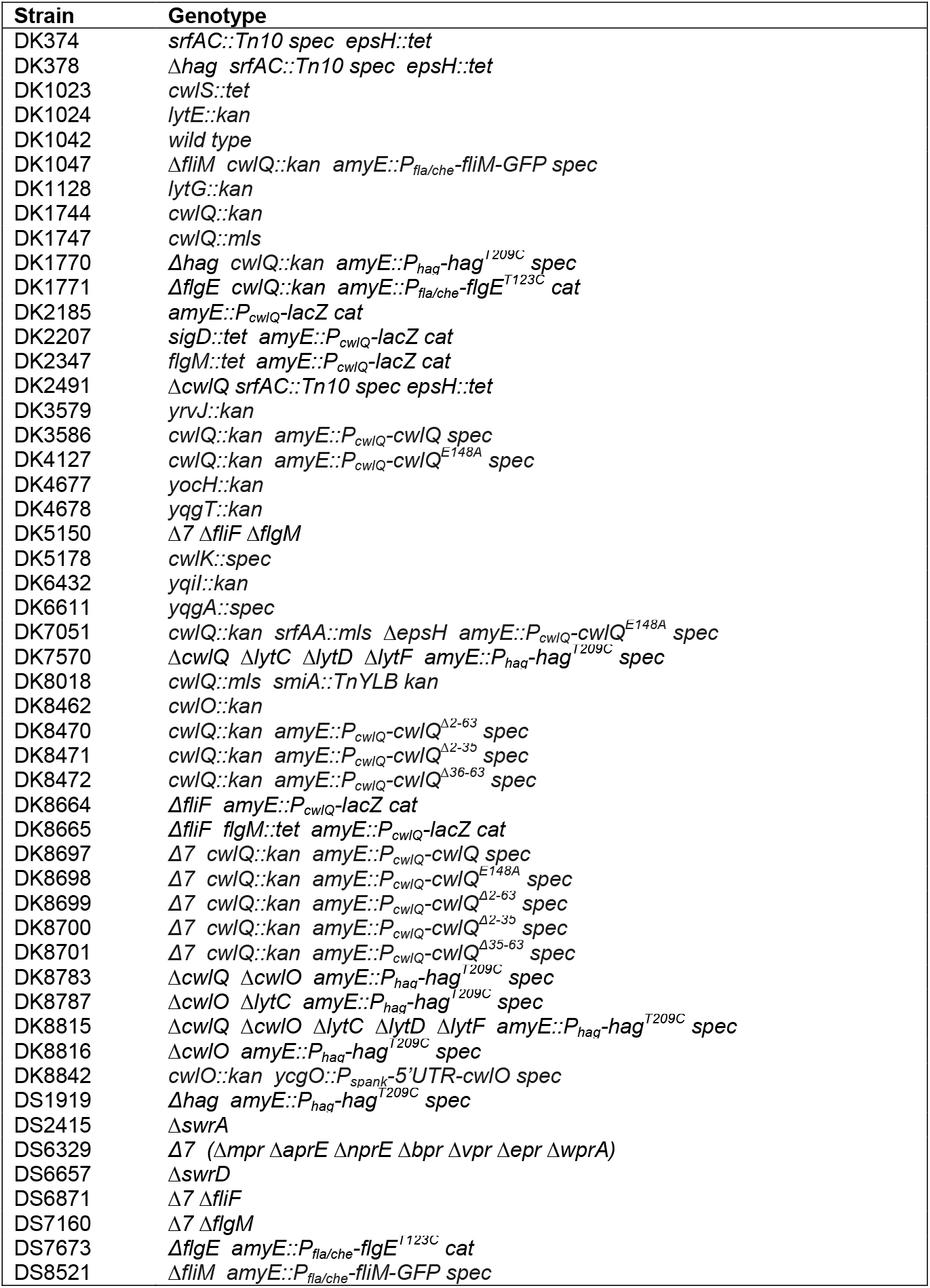
Strains

| Strain | Genotype |
| --- | --- |
| DK374 | *srfAC::Tn10 spec epsH::tet* |
| DK378 | Δ*hag srfAC::Tn10 spec epsH::tet* |
| DK1023 | *cwlS::tet* |
| DK1024 | *lytE::kan* |
| DK1042 | wild type |
| DK1047 | Δ*fliM cwlQ::kan amyE::P<sub>fla/che</sub>-fliM-GFP spec* |
| DK1128 | *lytG::kan* |
| DK1744 | *cwlQ::kan* |
| DK1747 | *cwlQ::mIs* |
| DK1770 | Δ*hag cwlQ::kan amyE::P<sub>haq</sub>-hag<sup>T209C</sup> spec* |
| DK1771 | Δ*flgE cwlQ::kan amyE::P<sub>fla/che</sub>-flgE<sup>T123C</sup> cat* |
| DK2185 | *amyE::P<sub>cwlQ</sub>-lacZ cat* |
| DK2207 | *sigD::tet amyE::P<sub>cwlQ</sub>-lacZ cat* |
| DK2347 | *flgM::tet amyE::P<sub>cwlQ</sub>-lacZ cat* |
| DK2491 | Δ*cwlQ srfAC::Tn10 spec epsH::tet* |
| DK3579 | *yrvJ::kan* |
| DK3586 | *cwlQ::kan amyE::P<sub>cwlQ</sub>-cwlQ spec* |
| DK4127 | *cwlQ::kan amyE::P<sub>cwlQ</sub>-cwlQ<sup>E148A</sup> spec* |
| DK4677 | *yocH::kan* |
| DK4678 | *yqgT::kan* |
| DK5150 | Δ7 Δ*fliF* Δ*flgM* |
| DK5178 | *cwlK::spec* |
| DK6432 | *yqil::kan* |
| DK6611 | *yqgA::spec* |
| DK7051 | *cwlQ::kan srfAA::mIs ΔepsH amyE::P<sub>cwlQ</sub>-cwlQ<sup>E148A</sup> spec* |
| DK7570 | Δ*cwlQ ΔlytC ΔlytD ΔlytF amyE::P<sub>haq</sub>-hag<sup>T209C</sup> spec* |
| DK8018 | *cwlQ::mIs smiA::TnYLB kan* |
| DK8462 | *cwlO::kan* |
| DK8470 | *cwlQ::kan amyE::P<sub>cwlQ</sub>-cwlQ<sup>Δ2-63</sup> spec* |
| DK8471 | *cwlQ::kan amyE::P<sub>cwlQ</sub>-cwlQ<sup>Δ2-35</sup> spec* |
| DK8472 | *cwlQ::kan amyE::P<sub>cwlQ</sub>-cwlQ<sup>Δ36-63</sup> spec* |
| DK8664 | Δ*fliF amyE::P<sub>cwlQ</sub>-lacZ cat* |
| DK8665 | Δ*fliF flgM::tet amyE::P<sub>cwlQ</sub>-lacZ cat* |
| DK8697 | Δ7 *cwlQ::kan amyE::P<sub>cwlQ</sub>-cwlQ spec* |
| DK8698 | Δ7 *cwlQ::kan amyE::P<sub>cwlQ</sub>-cwlQ<sup>E148A</sup> spec* |
| DK8699 | Δ7 *cwlQ::kan amyE::P<sub>cwlQ</sub>-cwlQ<sup>Δ2-63</sup> spec* |
| DK8700 | Δ7 *cwlQ::kan amyE::P<sub>cwlQ</sub>-cwlQ<sup>Δ2-35</sup> spec* |
| DK8701 | Δ7 *cwlQ::kan amyE::P<sub>cwlQ</sub>-cwlQ<sup>Δ35-63</sup> spec* |
| DK8783 | Δ*cwlQ ΔcwlO amyE::P<sub>haq</sub>-hag<sup>T209C</sup> spec* |
| DK8787 | Δ*cwlO ΔlytC amyE::P<sub>haq</sub>-hag<sup>T209C</sup> spec* |
| DK8815 | Δ*cwlQ ΔcwlO ΔlytC ΔlytD ΔlytF amyE::P<sub>haq</sub>-hag<sup>T209C</sup> spec* |
| DK8816 | Δ*cwlO amyE::P<sub>haq</sub>-hag<sup>T209C</sup> spec* |
| DK8842 | *cwlO::kan ycgO::P<sub>spank</sub>-5'UTR-cwlO spec* |
| DS1919 | Δ*hag amyE::P<sub>haq</sub>-hag<sup>T209C</sup> spec* |
| DS2415 | Δ*swrA* |
| DS6329 | Δ7 (Δ*mpr ΔaprE ΔnprE Δbpr Δvpr Δepr ΔwprA*) |
| DS6657 | Δ*swrD* |
| DS6871 | Δ7 Δ*fliF* |
| DS7160 | Δ7 Δ*flgM* |
| DS7673 | Δ*flgE amyE::P<sub>fla/che</sub>-flgE<sup>T123C</sup> cat* |
| DS8521 | Δ*fliM amyE::P<sub>fla/che</sub>-fliM-GFP spec* |

### Antibiotic resistance cassette insertion/deletion mutations

For each gene mutated, a PCR fragment was amplified upstream of the gene and downstream of the gene using the indicated primer pairs. Next, the antibiotic resistance cassette amplified from either pDG1515 (for tetracycline resistance), pDG780 (for kanamycin resistance) or pAH52 (for macrolide, lincomycin, streptomycin “mls” resistance) using primer pair 3250/3251 (72,73). Finally the three products were assembled by Gibson Isothermal Assembly (ITA) (74) and transformed into DK1042 by natural transformation. Finally, colonies containing the antibiotic resistance marker replacement mutant were determined by PCR amplification over the top the allele using the far upstream and far downstream primers used to generate the corresponding arms of adjacent DNA. The following primers were used to generate the indicated mutants: *cwlQ* (3695/3696::3697/3698); *cwlS* (3699/3700::3701/3702); *lytE* (3670/3671::3672/3673); *lytG* (3707/3708::3709/3710); *yqgT* (5497/5498::5499/5500); *yocH* (5501/5502::5503/5504); *yqgA* (5112/5113::5114/5115); *yqiL* (6425/6426::6427/6428); and *yrvJ* (4618/4619::4620/4621).

### *ΔcwlQ* in-frame markerless deletion

To generate the *ΔcwlQ* in frame marker-less deletion construct, the region upstream of *cwlQ* was PCR amplified using the primer pair 4118/4120 and the region downstream of *cwlQ* was PCR amplified using the primer pair 4119/4121. The two fragments were combined with SalI-digested pMiniMAD which carries a temperature sensitive origin of replication and an erythromycin resistance cassette (75) and were assembled by ITA to generate plasmid pSS5. The pSS5 plasmid was passaged though the *recA^+^ E. coli* strain TG1 before being transformed into DK1042 and selecting for *mls* resistance at the non-permissive temperature for plasmid replication, 37°C. To evict the plasmid, the strain was incubated in 3ml LB broth at a permissive temperature for plasmid replication (22°C) for 14 hours, diluted 30-fold in fresh LB broth, then serially diluted and plated on LB agar at 37°C. Individual colonies were patched on LB plates and LB plates containing *mls* to identify *mls* sensitive colonies that had evicted the plasmid. Chromosomal DNA from colonies that had excised the plasmid was purified and screened by PCR using primers 4118/4121 to determine which isolate had retained the *ΔcwlQ* allele.

### *ΔcwlO* markerless deletion

A *cwlO::kan* mutant allele generated from high throughput directed mutagenesis (76) was requested from the Bacillus Genetic Stock Center (The Ohio State University, Columbus OH). The kanamycin resistance cassettes is flanked by lox recombination sites and was excised by transformation with plasmid pDR244 encoding the *cre* recombinase and a spectinomcyin resistance cassette by plating on LB containing spectinomycin at 30°C. Colonies were restruck on LB, grown at 37°C and deletion of cwlO was determined by PCR product length polymorphism using primers 4741/4744.

### *cwlQ* complementation constructs

To generate the amyE::P*_cwlQ_-cwlQ* complementation construct (pSS9), a PCR product containing the *cwlQ* coding region plus 393 base pairs of upstream sequence was amplified from *B. subtilis* 3610 chromosomal DNA using the primer pair 4373/4374, digested with *BamHI* and *EcoRI* and cloned into the *BamHI* and *EcoRI* sites of pAH25 containing a polylinker and spectinomycin resistance cassette between two arms of the *amyE* (generous gift of Dr. Amy Camp, Mount Hoyloke College).

The active site mutant *amyE::P_cwlQ_-cwlQ^E148A^* allele construct was generated using a modified ITA protocol. Briefly, the region upstream of the complementation construct of *cwlQ* (DK3586) was PCR amplified using the primer pair 953/4888 and the region downstream of the complementation strain was PCR amplified using the primer pair 4887/954. The two fragments were assembled by isothermal assembly and retransformed into *B. subtilis* selecting for spectinomycin resistance. The N-terminal *cwlQ* deletion constructs were built using a similar approach with the indicated primer pair sets: *amyE::P_cwlQ_-cwlQ^Δ2-63^* (953/7304::7303/954), *amyE::P_cwlQ_-cwlQ^Δ2-35^* (953/7389::7388/954), and *amyE::P_cwlQ_-cwlQ^Δ36-63^* (953/7387::7386/954).

### *P_cwlQ_-lacZ* reporter construct

To generate the *P_cwlQ_-lacZ* reporter construct pSS2, the *P_cwlQ_* promoter was amplified from *B. subtilis* 3610 chromosomal DNA using the primers 4114/4115, digested with *Eco*RI and *Bam*HI and cloned into the *Eco*RI and *Bam*HI sites of plasmid pDG268, which carries a chloramphenicol-resistance marker and a polylinker upstream of the *lacZ* gene between two arms of the *amyE* gene (77).

### CwlQ-His expression construct and CwlQ purification

To generate the CwlQ-6His expression plasmid pSS14, the *cwlQ* gene was PCR amplified 3610 chromosomal DNA with primers 4757/4758, digested with EcoRI and HinDIII and cloned into the EcoRI and HinDIII sites of pET21a (Novagen). Next, pSS14 was transformed into Rosetta gami *E. coli*, grown to an OD_600_ of 0.6 in 1 liter of LB broth, induced with 1 mM IPTG, and grown for 16 h at 16°C. Cells were pelleted and resuspended at room temperature in lysis buffer (50 mM Tris [pH 8.0], 300 mM NaCl, 10% glycerol) and treated with lysozyme and DNase I, and the lysis was carried out using a pressurized cell homogenizer. The lysed cells were centrifuged at 15,000 rpm for 30 min. The cleared supernatants were combined with Ni-NTA resin (Novagen) and immediately poured onto a 1-cm separation column (Bio-Rad); the resin was allowed to pack and was washed with lysis buffer. CwlQ-6His bound to the resin was then eluted with elution buffer (50 mM Tris [pH 8.0], 300 mM NaCl, 10% glycerol, 100 mM imidazole). The elution fractions were then run on SDS-PAGE gels and appropriate fractions were then pooled and concentrated to 2 ml. The final purification of CwlQ-6His protein was conducted via size exclusion chromatography on a Superdex 75 16/60 (GE Healthcare) column using gel filtration buffer (25 mM Tris [pH 8.0], 300 mM NaCl, 10% glycerol) and submitted to Cocalico (Stephens, PA), for injection into rabbits and polyclonal antibody generation.

### Isothermal assembly reaction

First a 5X ITA stock mixture was generated (500 mM Tris-HCL (pH 7.5), 50 mM MgCl_2_, 50 mM DTT (Bio-Rad), 31.25 mM PEG-8000 (Fisher Scientific), 5.02 mM NAD (Sigma Aldrich), and 1 mM of each dNTP (New England BioLabs)), aliquoted and stored at −80° C. An assembly master mixture was made by combining prepared 5X isothermal assembly reaction buffer (131 mM Tris-HCl, 13.1 mM MgCl_2_, 13.1 mM DTT, 8.21 mM PEG-8000, 1.32 mM NAD, and 0.26 mM each dNTP) with Phusion DNA polymerase (New England BioLabs) (0.033 units/μL), T5 exonuclease diluted 1:5 with 5X reaction buffer (New England BioLabs) (0.01 units/μL), Taq DNA ligase (New England BioLabs) (5328 units/μL), and additional dNTPs (267 μM). The master mix was aliquoted as 15 μl and stored at −80°C. DNA fragments were combined at equimolar amounts to a total volume of 5 μL and added to a 15 μl aliquot of prepared master mix. The reaction was incubated for 60 minutes at 50° C.

### SPP1 phage transduction

To 0.1 ml of dense culture grown in TY broth (LB broth supplemented after autoclaving with 10 mM MgSO_4_ and 100 μM MnSO_4_), serial dilutions of SPP1 phage stock were added and statically incubated for 15 minutes at 37°C. To each mixture, 3 ml TYSA (molten TY supplemented with 0.5% agar) was added, poured atop fresh TY plates, and incubated at 37°C overnight. Top agar from the plate containing near confluent plaques was harvested by scraping into a 50 ml conical tube, vortexed, and centrifuged at 5,000 x g for 10 minutes. The supernatant was treated with 25 μg/ml DNase final concentration before being passed through a 0.45 μm syringe filter and stored at 4°C. Recipient cells were grown to stationary phase in 2 ml TY broth at 37°C. 0.9 ml cells were mixed with 5 μl of SPP1 donor phage stock. 9 ml of TY broth was added to the mixture and allowed to stand at 37°C for 30 minutes. The transduction mixture was then centrifuged at 5,000 x g for 10 minutes, the supernatant was discarded, and the pellet was resuspended in the remaining volume. 100 μl of cell suspension was then plated on TY fortified with 1.5% agar, the appropriate antibiotic, and 10 mM sodium citrate.

### Motility assays

For the swarm expansion assay, cells were grown to mid-log phase at 37°C in LB broth and resuspended to 10 OD_600_ in pH 8.0 PBS buffer (137 mM NaCl, 2.7 mM KCl, 10 mM Na_2_HPO_4_, and 2 mM KH_2_PO_4_) containing 0.5% India ink (Higgins). Freshly prepared LB containing 0.7% Bacto agar (25 ml/plate) (For percent agar concentration assay the freshly prepared LB containing between 0.5% to 0.9%) was dried for 10 minutes in a laminar flow hood, centrally inoculated with 10 μl of the cell suspension, dried for another 10 minutes, and incubated at 37°C. The India ink demarks the origin of the colony and the swarm radius was measured relative to the origin. For consistency, an axis was drawn on the back of the plate and swarm radii measurements were taken along this transect.

For swim assays, cells were grown to mid-log phase at 37°C in LB broth and resuspended to 10 OD_600_ in pH 8.0 PBS buffer (137 mM NaCl, 2.7 mM KCl, 10 mM Na_2_HPO_4_, and 2 mM KH_2_PO_4_) 10 ul of culture were inoculated into the agar. Freshly prepared LB containing 0.3% Bacto agar (25 ml/plate) was dried for 10 minutes in a laminar flow hood, centrally inoculated with 10 μl of the cell suspension, dried for another 10 minutes, and incubated at 37°C. Plates were visualized with a BioRad Geldoc system and digitally captured using BioRad Quantity One software.

### Western blotting

*B. subtilis* strains were grown in LB broth to OD_600_ ~0.5, 10 ml was harvested by centrifugation, and resuspended to 100 OD_600_ in Lysis buffer (20 mM Tris pH 7.0, 10 mM EDTA, 1 mg/ml lysozyme, 10 μg/ml DNAse I, 100 μg/ml RNAse I, 1 mM PMSF) and incubated 30 minutes at 37°C. Each lysate was then mixed with the appropriate amount of 6x SDS loading dye to dilute the loading dye to 1x concentration. Samples were separated by 12% Sodium dodecyl sulfate-polyacrylamide gel electrophoresis (SDS-PAGE). The proteins were electroblotted onto nitrocellulose and developed with a 1:1,000 dilution of (anti-CwlQ) or 1:80,000 dilution of (anti-SigA) of primary antibody and a 1:10,000 dilution secondary antibody (horseradish peroxidase-conjugated goat anti-rabbit immunoglobulin G). Immunoblot was developed using the Immun-Star HRP developer kit (Bio-Rad).

For experiments involving trichloroacetic acid (TCA) precipitation, the 10 ml of supernatant was saved during pelleting step, combined with 1 ml 0.015% sodium deoxycholate, votexted and incubated 10 minutes at room temperature. Next, 500 ul of ice cold TCA was added, the mixture was vortexed and incubated on ice for 2 hours. The supernatant was precipitated by centrifugation at 30,000 x g for 10 minutes at 4°C. The pellet was resuspended in 1 ml ice cold acetone and repelleted in a tabletop centrifuge. Finally the pellet was resuspended in the same amount of 1X protein sample buffer as the corresponding pellet.

### Microscopy

Fluorescence microscopy was performed with a Nikon 80i microscope along with a phase contrast objective Nikon Plan Apo 100X and an Excite 120 metal halide lamp. Alexa Fluor 594 C_5_ maleimide fluorescent signals were visualized with a C-FL HYQ Texas Red Filter Cube (excitation filter 532-587 nm, barrier filter >590 nm). GFP was visualized using a C-FL HYQ FITC Filter Cube (FITC, excitation filter 460-500 nm, barrier filter 515-550 nm). YFP was visualized using a C_FL HYQ YFP Filter Cube (excitation filter 490-510 nm, barrier filter 515-550 nm). TMA-DPH fluorescent signal was visualized using a UB-2E/C DAPI Filter Cube (excitation filter 340-380 nm, barrier filter 435-485 nm). Images were captured with a Photometrics Coolsnap HQ^2^ camera in black and white, false colored and superimposed using Metamorph image software.

For super-resolution microscopy using structured illumination the OMX 3D-SIM Super Resolution system at Indiana University Bloomington Light Microscopy Imaging Center was used. Super-resolution microscopy was performed using a 1.4NA Olympus 100X oil objective. FM4-64 was visualized using laser line 561nm and emission filter 609-654nm, and Alexa Fluor 488 was visualized using laser line 488nm and emission filter 500-550nm. Images were captured by Photometrics Cascade II EMCCD camera and processed by SoftWoRx imaging software (Applied Precision). For counting hooks, images reconstructed with SoftWoRx were used in Imaris (Bitplane) to determine the number of FlgE^T123C^ foci on the surface of each cell. The spots feature labelled each FlgE^T123C^ foci by the search parameter of identifying spots of 1 μM in the 488 wavelength and we verified by eye that the spots labelling identified bonafide foci on the cell surface. The hook count datasets were plotted against cell length (micron) using IMARIS software.

## ACKNOWLEDGEMENTS

We thank Felix Dempwolff for intellectual and technical support, as well as strain construction. The work was funded by the National Institutes of Health R35 grant GM131783 to DBK.

---

**Figure S1. Most mutants in PG hydrolase candidates were wild type for swarming motility.** Quantitative swarm expansion assay of wild type (open circles) and the indicated mutant (closed circles). Each data point is the average of three replicates. The same wild type data may be repeated for multiple panels as one wild type control set was performed in swarm expansion assays with the corresponding mutants on the same day. The following strains were used to generate the panels: wild type (DK1042), *cwlK* (DK5178), *lytE* (DK1024), *lytG* (DK1128), *cwlS* (DK1023), *yocH* (DK4677), *yqgA* (DK6611), *yqgT* (DK4678), *yqiI* (DK6423), and *yrvJ* (DK3579).

